# Memory consolidation in honey bees is enhanced by down-regulation of *Down Syndrome Cell Adhesion Molecule* and changes its alternative splicing

**DOI:** 10.1101/2023.10.19.563081

**Authors:** Pinar Ustaoglu, David W. J. McQuarrie, Anthony Rochet, Thomas Dix, Irmgard U. Haussmann, Roland Arnold, Jean-Marc Devaud, Matthias Soller

## Abstract

Down syndrome cell adhesion molecule (*Dscam*) gene encodes a cell adhesion molecule required for neuronal wiring. A remarkable feature of invertebrate *Dscam* is massive alternative splicing generating thousands of different isoforms from three variable clusters of alternative exons. *Dscam* expression and diversity arising from alternative splicing have been studied during development, but whether they exert functions in differentiated brains has not been determined. Here, using honey bees, we find that Dscam expression is critically linked to memory retention as reducing expression by RNAi enhances memory after reward learning in adult worker bees. Moreover, alternative splicing of *Dscam* is altered in all three variable clusters after learning. Since identical Dscam isoforms engage in homophilic interactions, these results suggest a mechanism to alter inclusion of variable exons during memory consolidation to modify neuronal connections for memory retention.

## Introduction

Memory consolidation is the process through which memories are stabilized and stored into long-term memory (Kandel et al., 2014). Gene expression through transcription of genes plays pivotal roles in this process (Alberini, 2009, Alberini et al., 1994, Clayton, 2000, Kandel et al., 2014). During transcription, splicing out of introns allows to alter coding content through alternative splicing, which is particularly abundant in the brain in genes coding for ion channels and cell adhesion molecules (Soller, 2006, Vuong et al., 2016, Ule and Blencowe, 2019). From some of these genes, hundreds of different proteins can be made leading to a substantial increase of the coding capacity of the genome. However, the functional consequences of diversity generated by alternative splicing remain poorly understood.

In honey bees, an invertebrate model species for the study of learning and memory processes, requirement of transcription and the neuronal RNA binding protein ELAV during memory consolidation further suggest important roles for alternative splicing (Lefer et al., 2012, Villar et al., 2020, Ustaoglu et al., 2021). Accordingly, alternative splicing events in a number of cell adhesion molecules and ion channels among other genes have been linked to learning and memory (Demares et al., 2014, Demares et al., 2013, Beffert et al., 2005, Poplawski et al., 2016, Sengar et al., 2019, Ustaoglu et al., 2021). Likewise, ELAV itself is substantially alternatively spliced in bee mushroom bodies (brain centers for learning and memory) and its expression pattern differs between individuals, likely reflecting differences in individual experiences (Ustaoglu et al., 2021).

The *Down syndrome cell adhesion molecule* (*Dscam*) gene from invertebrates is the most extreme example for alternative splicing generating a mere 36’012 different proteins from a single gene in *Drosophila* and this likely is still an underestimate as new exons are still discovered (Hemani and Soller, 2012, Zhang et al., 2019). The main part of diversity in the *Dscam* gene originates from three clusters containing an array of variable exons of which only one exon is included. These clusters are in exon 4 with 12 variables, exon 6 with 48 variables and exon 9 with 36 variables in the extracellular domain and in exon 11 in the transmembrane domain with two variables (Schmucker et al., 2000).

*Drosophila Dscam* is expressed in the nervous system where its variability is used during development for neuronal wiring, and in the immune system as a pattern recognition receptor for pathogen clearance by phagocytosis (Hemani and Soller, 2012, Watson et al., 2005). The main roles in the nervous system comprise establishment of overlapping dendritic fields in the peripheral nervous system and to branch neurites in neuronal tracts (Hughes et al., 2007, Matthews et al., 2007, Soba et al., 2007). Key to these functions of Dscam are homophilic repulsive properties of individual identical Dscam isoforms, but not to other splice variants (Wojtowicz et al., 2004, Wojtowicz et al., 2007). Accordingly, dendrites of neurons can overlap if they express different Dscam isoforms and neuronal tracts such as the *Drosophila* mushroom body containing hundreds of neurons can split into two lobes (Wang et al., 2002, Matthews et al., 2007).

Mechanistically, it has been proposed that individual neurons adopt a unique set of variables stochastically to comply with their fate in branching in the peripheral nervous system and in the mushroom bodies (Neves et al., 2004, Miura et al., 2013). This model further implies that *Dscam* alternative splicing is maintained in individual neurons. In the immune system, however, exposure to different pathogens altered inclusion of variables to generate variants with higher affinity in mosquitoes (Dong et al., 2006). Accordingly, both stochastic as well as deterministic rules have been found to govern dendritic branching (Palavalli et al., 2021). Moreover, the homophilic interaction of identical Dscam isoforms could also be converted to attraction at low levels and in this way build novel connections (Wojtowicz et al., 2004).

Although vertebrate *Dscam* does not have the molecular diversity generated by alternative splicing as found in invertebrates, the homophilic properties of Dscam are a conserved molecular function required for neuronal wiring (Schmucker and Chen, 2009). Since the *Dscam* gene in humans has been associated with intellectual disability (Vacca et al., 2019), we examined its role in learning and memory in honey bees, where memory consolidation involves changes in gene expression (Lefer et al., 2012, Villar et al., 2020, Ustaoglu et al., 2021) and neuronal connectivity (Hourcade et al., 2010). Here, we discovered that down-regulated *Dscam* during learning enhances memory consolidation in bees. To characterize alternative splicing in bees we sequenced *Dscam* cDNAs of the variable regions to comprehensively update previous annotation from comparing genomic sequences (Graveley et al., 2004). This analysis reveals that bees have 9 exon 4, 52 exon 6 and 19 exon 10 variables. Unlike *Drosophila*, we find intra-cluster splicing in bee *Dscam*. When we analysed alternative splicing during the course of memory consolidation, we discovered that alternative splicing significantly changed within variable clusters. Our results indicate that changes in *Dscam* splicing can be induced upon experience likely to establish new neuronal connections associated with memory consolidation.

## Results

### Dscam is required for memory consolidation in bees after olfactory reward conditioning

To detect bee Dscam, we used a polyclonal anti-serum raised against *Drosophila* Dscam (Watson et al., 2005). In honey bees this polyclonal antibody recognizes proteins at the size of *Drosophila* Dscam of 222 and 270 kDa and three shorter bands of 100, 110 and 130 kDa (Supplementary Fig 1A). To validate that the larger band recognized by the antibody is indeed bee Dscam, we injected dsRNA against bee *Dscam* into the central brain for knock-down by RNAi. A substantial reduction of 72±5% and 79±10% (n=3 each) is achieved after 48 h and 64 h of the expected bands at 222 and 270 kDa, respectively, but not of the shorter bands of 100, 110 and 130 kDa suggesting that these bands in bees are unspecific (Fig 1A and Supplementary Fig 1B).

**Fig. 1.**
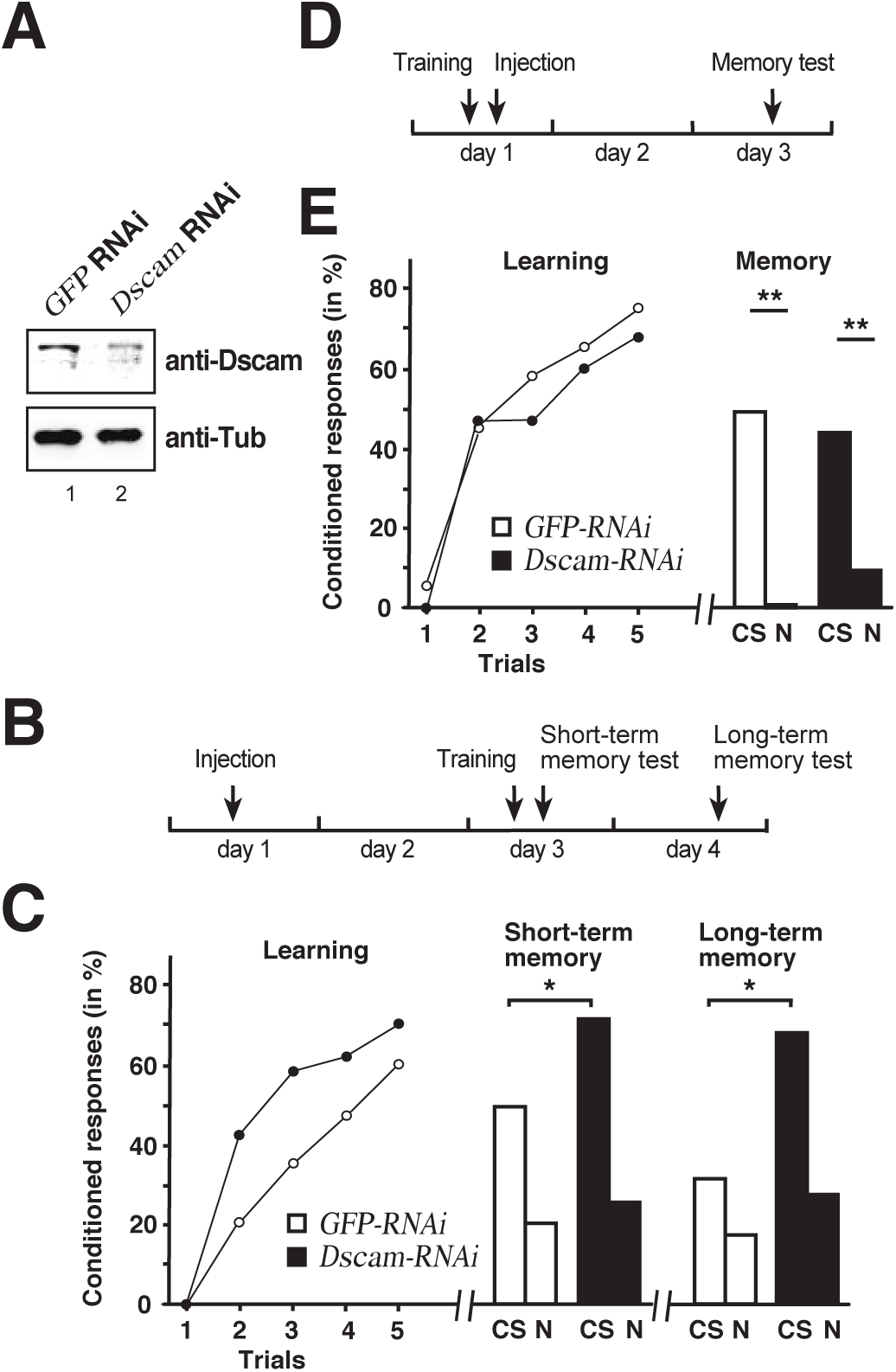
Dscam is required for learning and memory consolidation. **A** Western blot detecting Dscam (top) or tubulin (bottom) in bee central brains of control *GFP* and *Dscam* dsRNA injected workers 48 h after injection. **B** Schematic of the treatment to test for Dscam’s role in learning and memory consolidation. **C** Learning (*left*), short term (*middle*) and long-term memory (*right*) performances of control *GFP* dsRNA (*white*) and *Dscam* dsRNA (*black*) injected worker bees. *CS*: conditioned odor, *N*: novel odorant. *: p<0.05. **D** Schematic of the treatment control for Dscam’s role in learning and memory consolidation. **E** Learning (*left*) and memory (*right*) performances of control *GFP* dsRNA (*white*, n=39/n=22 at 24h) and *Dscam* dsRNA (*black*, n=46/n=15 at 24h) injected worker bees. *CS*: conditioned stimulus, *N*: novel odorant **: p<0.01. The source date underlying this figure are available in Supplementary Data 1.

To assess whether Dscam has a role in learning and memory, bees were individually trained in an associative olfactory conditioning task two days after injection of *Dscam* or *GFP* control dsRNA to then measure short-term and long-term memory 1 hour and 1 day after training, respectively (Fig 1B). Both groups showed significant learning over the successive trials (RM-ANOVA, *Trial* effect: F=46.55, p<0.001), without significant effect of the treatment (F=2.28, p=0.135, Fig 1C left). We then tested short-term (1h following conditioning) and long-term (24h) memory to ask whether *Dscam* downregulation could impact on memory formation despite preserved learning capacities (Fig 1C). Here, at 1 hour (Fig 1C middle) bees from the *Dscam* group responded significantly more to the conditioned stimulus (CS) than those from the *GFP* group (Fischer’s exact test: χ^2^ = 4.72, p=0.044), while the proportions of specific responses did not differ either (χ^2^ = 3.37, p=0.118, Fig 1C middle and left). Yet, while both groups clearly responded more to the CS than to the novel odorant 1 hour after training, the difference was not significant in either case (McNemar test, *GFP*: χ^2^ = 2.88, p=0.090 : *Dscam*: χ^2^ = 0.0588, p=0.808). The proportions of individuals displaying specific responses (i.e. responses to the CS but not the novel odour, N) did not differ either between the groups (χ^2^ = 2.56, p=0.125, Fig 1C middle). Thus, down-regulated Dscam levels enhanced memory performance without affecting its specificity.

At 24 hours after conditioning, only a portion of bees survived and could be tested (*GFP*: n=22, *Dscam*: n=15). Both groups responded more to the CS than to the novel odorant, although the difference was not significant in the *Dscam* group due to a smaller sample size (McNemar test, *GFP*: χ^2^ = 4.84, p=0.028; *Dscam*: χ^2^ = 0.048, p=0.827, Fig 1C right). Again, the proportions of specific responses did not differ between the groups (χ^2^ = 0.3.37, p=0.118), but bees from the Dscam group showed a marginally significantly higher response rate to the CS than those form the GFP group (Fischer’s exact test: χ^2^ = 4.36, p=0.050). Thus, down-regulated Dscam levels enhanced CS-N specific long-term memory performance.

To reject the possibility that loss of *Dscam* impacts on long-term memory retrieval per se due to a prolonged downregulation of *Dscam*, we performed an additional experiment in which injection was done shortly after training, so that RNAi would be effective at the time of retrieval (24 hours) rather than at consolidation (Fig 1D). As expected, this treatment did not affect learning (*Trial effect*: F=48.48, p<0.001; *Trial x Treatment* interaction: F=0.697, p>0.05; Fig 1E left). More importantly, long-memory retrieval was intact and two days after training both groups responded similarly to the CS (Fischer’s test: χ^2^ = 0.729, p>0.05) and responded significantly less to the novel odorant (GFP: χ^2^ = 5.23, p<0.05; elavl2: χ^2^ = 7.47, p<0.01), thus indicating a preserved memory of the CS-US association and no enhancement of memory (Fig 1E right).

These results thus argue that down-regulation of *Dscam* during memory consolidation enhances storage of memories.

### Alternative splicing analysis of honey bee *Dscam* reveals novel isoforms and intra-cluster splicing

Since *Dscam* in invertebrates is abundantly alternatively spliced, we wanted to analyze Dscam alternative splicing after training, but this required evaluation of the gene annotation. The exon-intron structure of honey bee *Dscam* had previously been annotated based phylogenomic conservation and comparison to cDNAs obtained from *Drosophila Dscam* (Graveley et al., 2004). To directly address alternative splicing in *Dscam* variable clusters 4, 6 and 10 in bees, we RT-PCR amplified the variable clusters and Illumina sequenced the amplicons. This analysis revealed strong overlap with the previous annotation, but also novel isoforms and additional intra-cluster splicing (Decio et al., 2019).

In particular, in each of the clusters we find an additional exon (exon 4.0, 6.0 and 10.0) before the previously annotated isoforms (Fig 2A-C). Exon 4.0 and 10.0 alter the frame resulting in non-productive isoforms.

**Fig. 2.**
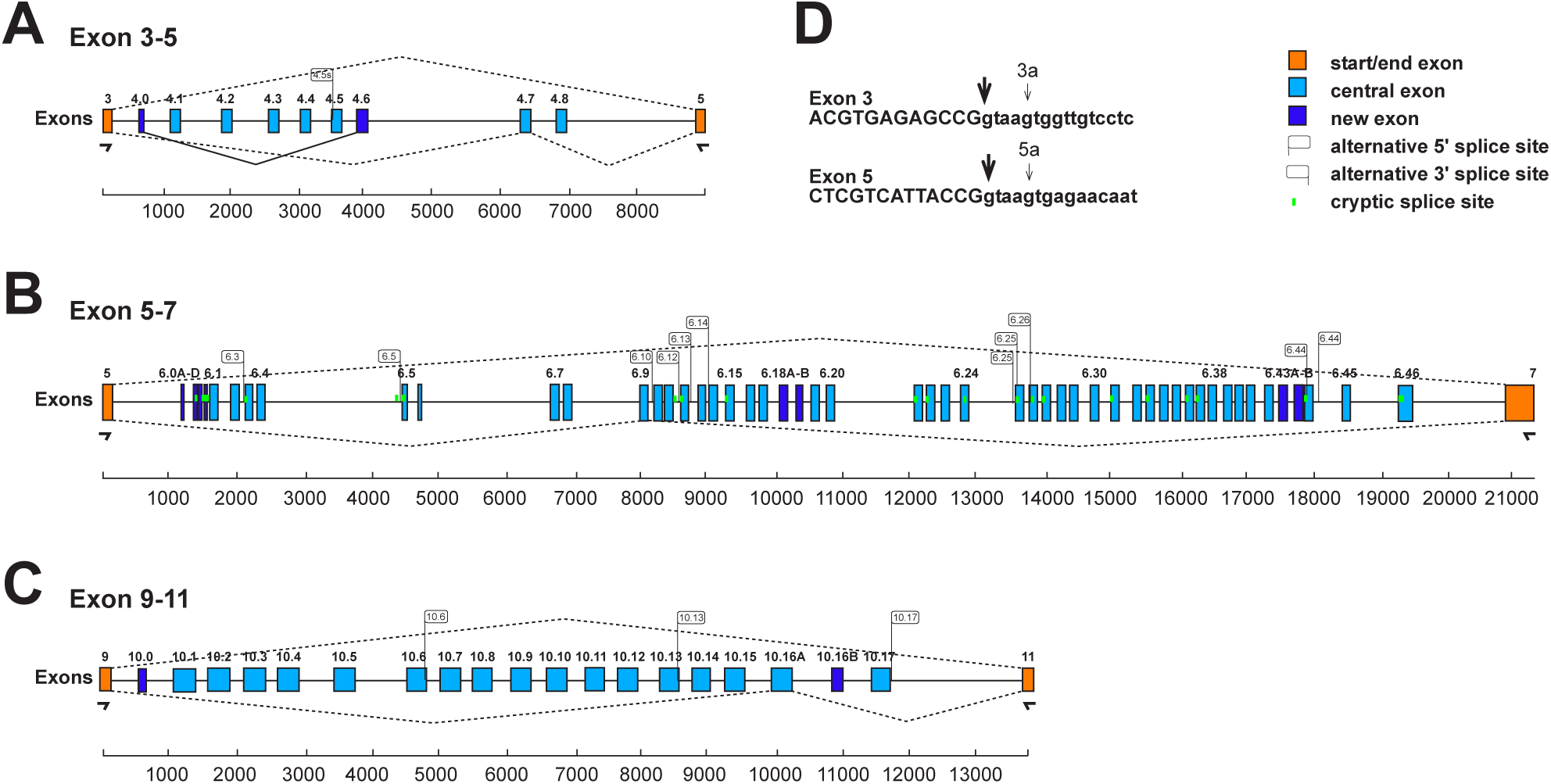
Schematic of the honey bee *Dscam* variable exon clusters 4, 6 and 10. **A-C** Gene models of *Apis mellifera Dscam* variable exon clusters 4 (A, top), 6 (B, middle) and 10 (C, bottom) depicting constant flanking exons in orange and variable exons in light blue boxes. New variable exons are indicated by dark blue boxes. Alternative 5′ and 3′ splice sites are indicated by right and left directed flags, respectively. Intra-cluster splicing is indicated by solid lines and dashed lines indicate representative alternative splicing. Cryptic splice sites (< 1%) are indicated by vertical green lines. Exon clusters are drawn to scale with scale bars are shown at the bottom. **D** Alternative 5’ splice sites in exons 3 and 5 are shown as small arrows.

In the exon 4 cluster, we find prominent use of an alternative 5′ splice sites in exon 3 that splices to exons 4.1 and 4.3 (3a4.1 and 3a4.3, Fig 2D) that changes the open reading frame to generate non-function products. In addition, an addition 3’ splice site is present in exon 4.5 generating an in-frame shorter isoform (Fig 2A).

In the beginning of the exon 6 cluster, we find a number of 3′ splice sites, which we describe as exons 6.0A-D, but we could not detect a downstream 5′ splice site by amplicon sequencing(Fig 2B). Potentially, they could include large exons using known 5′ splice sites.

In the exon 6 and 10 clusters (Fig 2B and 2C), we identified additional variable exons (6.18B, 6.43B and 10.16B), that maintain the reading frame and additional 5′ and 3′ splice sites, that are above splicing noise (> 1%). In the exon 6 cluster, we find one additional 5′ splice site (6.44s2) and 13 new 3′ splice sites (exons 6.1s, 6.3s, 6.5s, 6.10s, 6.11s, 6.12s, 6.13s, 6.14s, 6.15s, 6.25s1, 6.25s2, 6.26s, 6.31s, 6.33s, and 6.44s). In the exon 10 cluster, we find three additional 5′ splice sites (exon 10.6s, 10.13s and 10.17s). All of these new variable exons maintain the reading frame.

In addition, we find low levels of skipping of exons in the variable clusters. Skipping of exon 4 and 10 variables maintains the reading frame, while skipping of exon 6 variables leads to truncated Dscam isoforms.

### Alternative splicing in *Dscam* exon 4 cluster changes in slow learners during memory consolidation

To analyse alternative splicing of *Dscam* in the variable cluster 4, we used our previously established gel-based assay, that can distinguish all eight variables and exon skipping in addition to newly discovered isoforms and reliably detect changes during development and adult morphs (Haussmann et al., 2019, Decio et al., 2019). When we analysed inclusion of exon 4 variables we noticed obvious differences in some bees, particularly ones that have learned faster than others (Ustaoglu et al., 2021) and we therefore split trained bees into two groups of fast and slow learners. As negative controls, we used a group of unpaired bees, which were presented the stimuli (CS: odorant, US: sucrose) as trained bees, except that the delay between the CS and US was increased to impede the learning of any association.

Upon learning differences appeared immediately particularly for exons 4.1s and 4.3s as well as exon skipping, but only in some bees (Fig 3A). When we quantified the inclusion levels (Fig 3B-D), significant changes were observed in exons 4.0 and 4.6s two hours after training in the slow learners. Four hours after training, significant changes were detected for exons 4.2, 4.3, 4.6s and skipped exons in slow learners (Fig 3D).

**Fig. 3.**
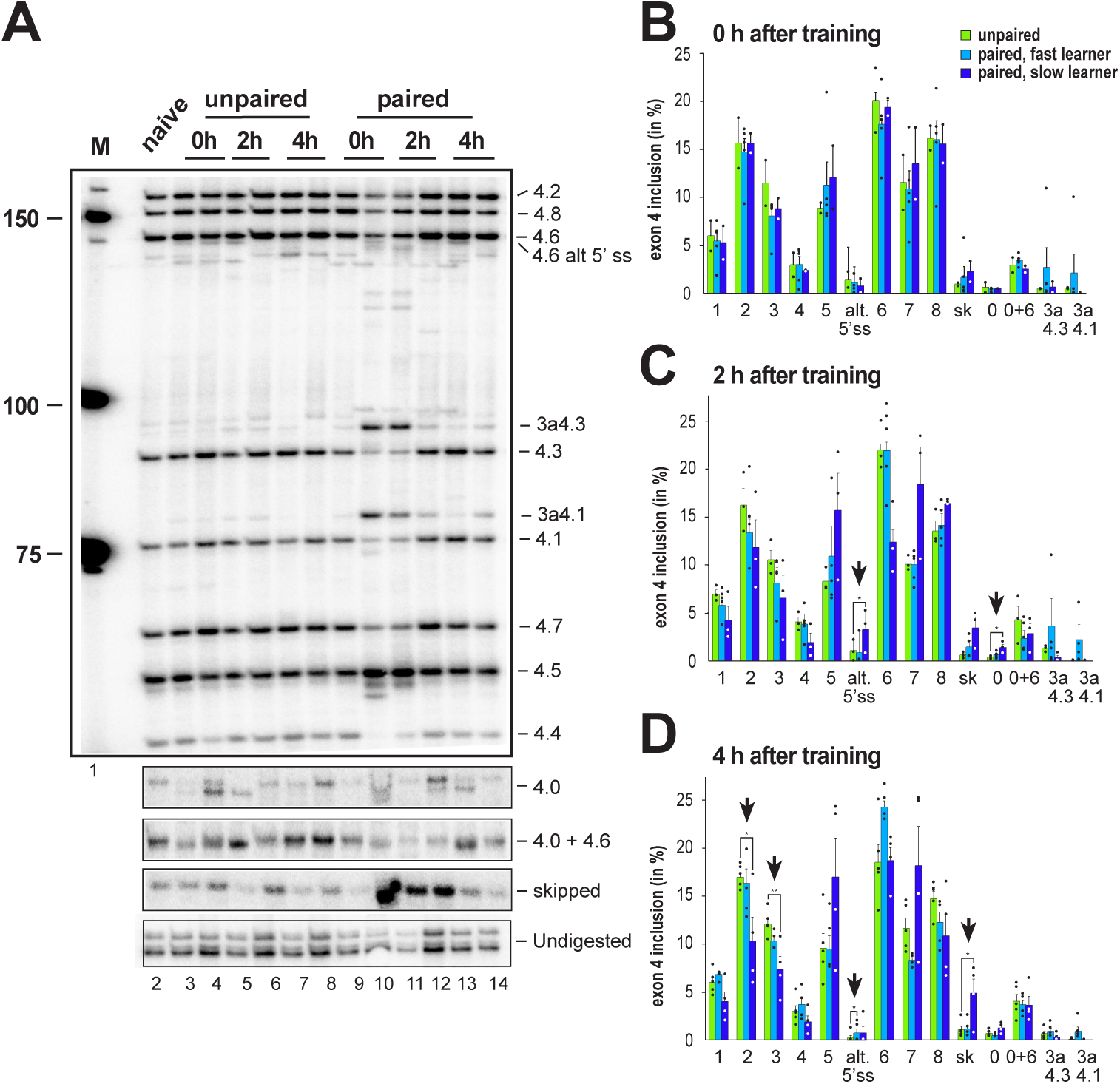
*Dscam* variable exon 4 alternative splicing analysis during memory consolidation. **A** Representative 5% denaturing polyacrylamide gel separating ^32^P-labeled alternative splice products from worker brains conditioned with unpaired (left) and paired odors and reward. Identity of splice variants are indicated at the right. M: Marker. **B-D** Analysis of alternative splicing unpaired control (green), fast (light blue) and slow learners (dark blue) indicating exon inclusion frequency (in %) for the different variables immediately after training (0 h), 2 h or 4 h after training. Significant changes are indicated by vertical arrows (*: p<0.05; **: p<0.01). The source date underlying this figure are available in Supplementary Data 1.

### Alternative splicing in Dscam exon 9 variables changes most prominently in slow learners two hours after training

To analyse alternative splicing of *Dscam* in the variable cluster 10, we also used our previously established gel-based assay, that can distinguish 13 from 18 variables (72%) and exon skipping with significant changes during development and in adult morphs (Fig 4A and Suppl. Fig 2).

**Fig. 4.**
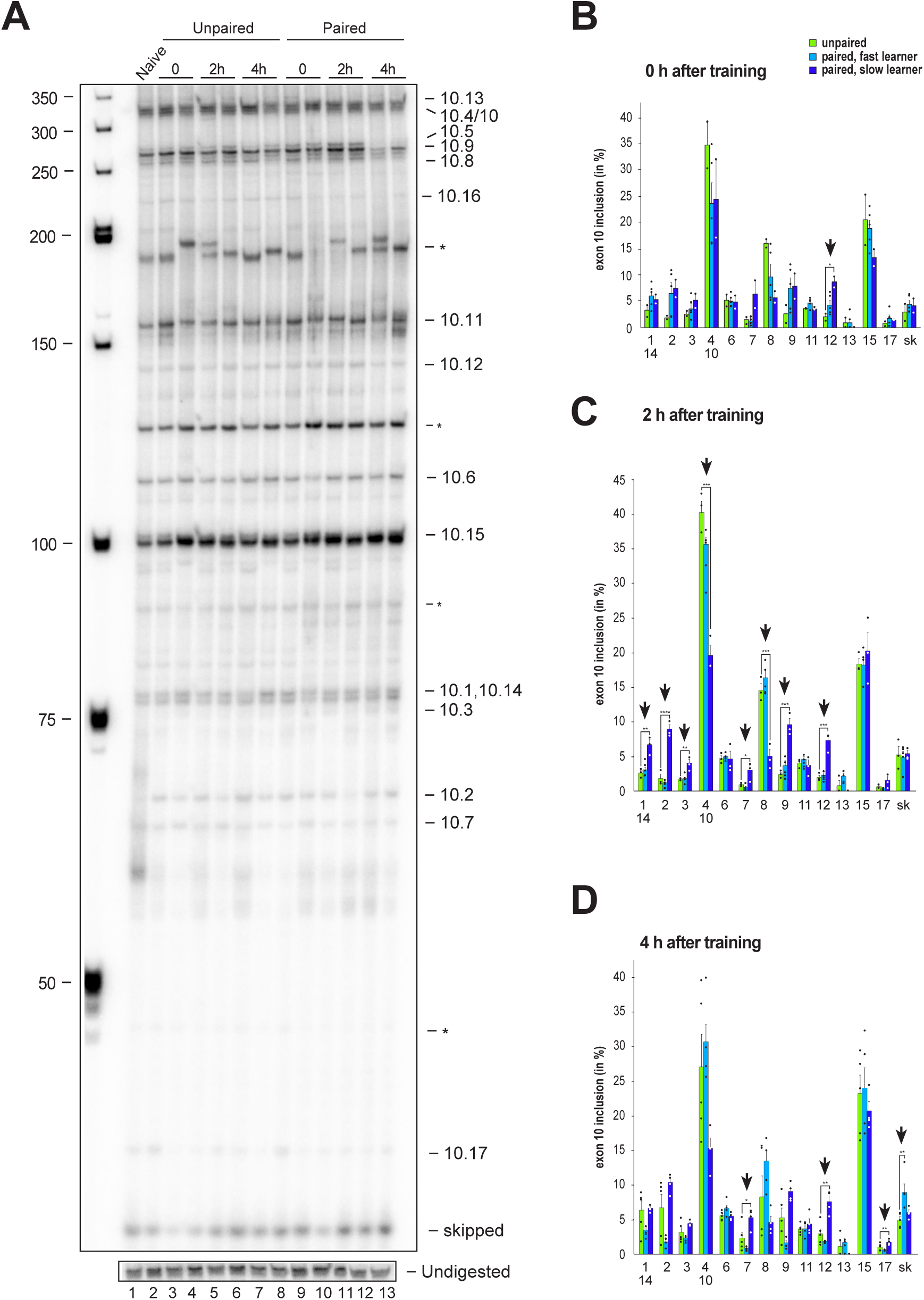
*Dscam* variable exon 10 alternative splicing analysis during memory consolidation. **A** Representative 5 % denaturing polyacrylamide gel separating ^32^P-labeled alternative splice products from worker brains conditioned with unpaired (left) and paired odors and reward. Identity of splice variants are indicated at the right. M: Marker. **B-D** Analysis of alternative splicing unpaired control (green), fast (light blue) and slow learners (dark blue) indicating exon inclusion frequency (in %) for the different variables immediately after training (0 h), 2 h or 4 h after training. Significant changes are indicated by vertical arrows (*: p<0.05; **: p<0.01). The source date underlying this figure are available in Supplementary Data 1.

Upon olfactory conditioning of bees, significant changes were detected immediately after training in slow learners for exon 10.12 (Fig 4B). Intriguingly, two hours after training significant changes were detected in 62 % (8 from 13) of variable exons in slow learners (Fig 4C). Four hours after training, significant changes were found in exons 10.7, 10.12. and 10.17 of slow learners, while fast learners show a significant increase in exon skipping (Fig 4D).

### Alternative splicing in *Dscam* of more than half of exon 6 variables changes in fast learners two hours after training

For the analysis of *Dscam* alternative splicing in the exon 6 cluster, we switched to amplicon sequencing as the gel-based assay could not sufficiently resolve the variables. For this analysis we did paired-end sequencing two hours after training and summed up the reads for control bees where no paired stimulus was given, and for bees that either learned fast or slow.

Strikingly, from the 48 different exon 6 variables, 63% (30 out of 48), showed significant changes in fast learning bees compared to controls, while no significant differences were detected between controls and slow learners (Fig 5A and 5B).

**Fig. 5.**
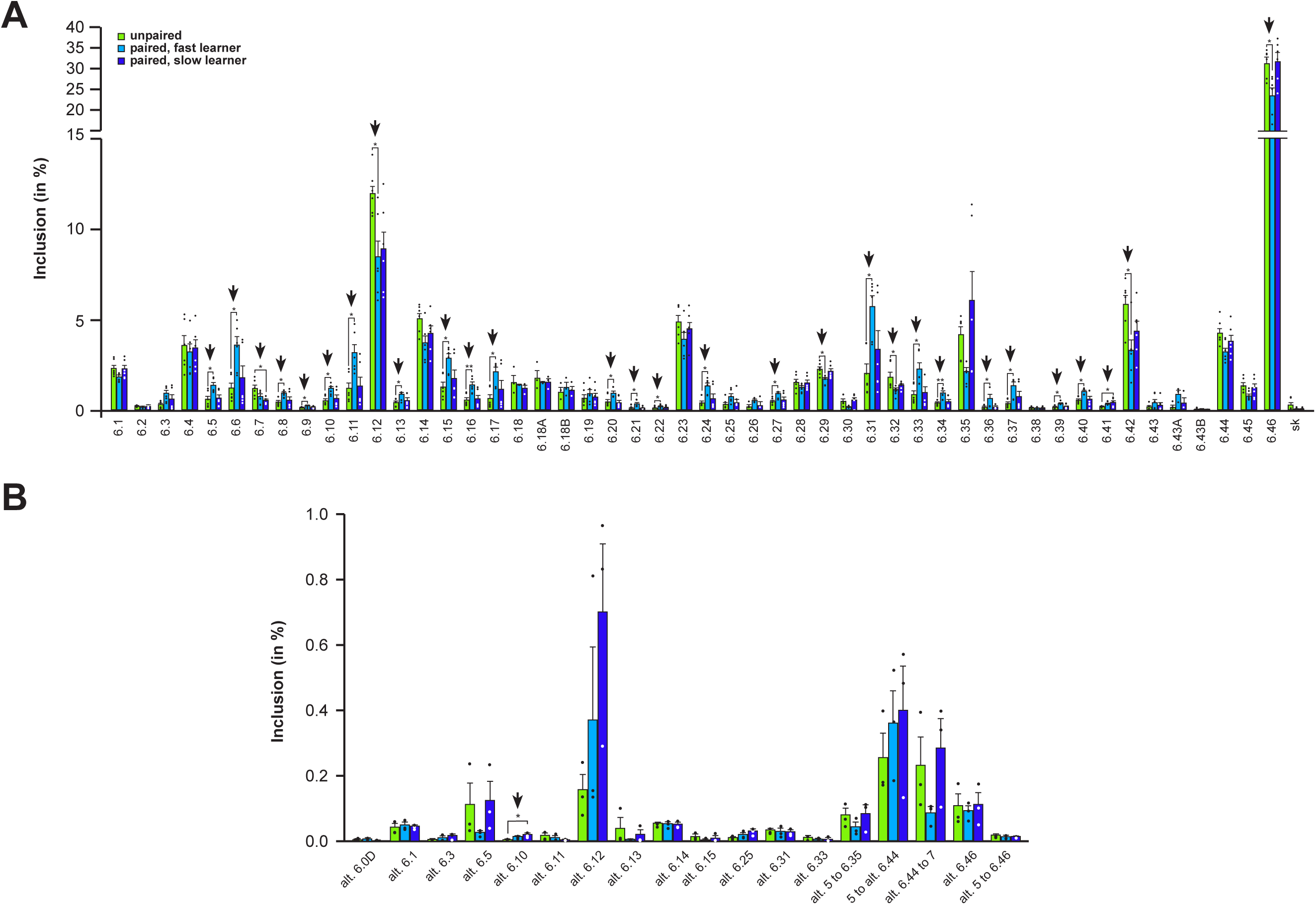
*Dscam* variable exon 6 alternative splicing analysis during memory consolidation. **A and B** Analysis of alternative splicing unpaired control (green), fast (light blue) and slow learners (dark blue) indicating exon inclusion frequency (in %) from amplicon sequencing for the different variables 2 h after training for main variable (A) and alternative splice sites (B). Significant changes are indicated by vertical arrows (*: p<0.05; **: p<0.01). The source date underlying this figure are available in Supplementary Data 1.

## Discussion

Alternative splicing massively expands proteomic diversity and is particularly abundant in the brain. It is most prominently elaborated in the invertebrate *Dscam* gene making about three times as many proteins than genes present in the genome (Hemani and Soller, 2012, Soller, 2006). Here, we characterize in depth the alternative splicing landscape of the honey bee *Dscam* gene and identify novel isoforms including an increased use of alternative 5′ and 3′ splice sites and intra-cluster splicing further expanding the diversity and shedding light on its unusual splicing regulation. *Dscam* mis-regulation has been associated with intellectual disabilities in humans associated with an additional copy of chromosome 21 (Vacca et al., 2019). In particular, an extra copy of *Dscam* in *Drosophila* causes strengthening of synapses and in a mouse model leads to excessive GABAergic synapses in the neocortex (Liu et al., 2023, Lowe et al., 2018). When we analysed learning and memory in honey bees upon reducing Dscam levels by RNAi we find enhanced memory storage. In addition, we find that alternative splicing of *Dscam* changes in the variable clusters during memory consolidation. Since Dscam homophilic interaction could stabilize existing connections (Chen et al., 2006), down-regulation of specific isoforms could loosen existing neuronal connections to stimulate formation of new ones, essentially recapitulating down-regulation of *Dscam* as obtained by RNA interference. To store individual memory traces in the mushroom bodies, likely only few neurons are involved. It is thus surprising that Dscam alternative splicing changed substantially, such that significant changes could be detected in central brains. It will be of interest to follow up which neurons exactly change splicing upon learning, which might be possible with the development of long-read sequencing technology for single cell analysis to overcome current limitations in alternative splicing analysis (Decio et al., 2021, Decio et al., 2019, Zaharieva et al., 2012).

Memory consolidation requires transcription and implicates that alternative splicing can impact on the process of memory storage (Kandel et al., 2014, Ustaoglu et al., 2021). In honey bees, a first transcriptional wave occurs at the beginning of memory consolidation (Lefer et al., 2012). Of note, it is during this phase that also *Dscam* alternative splicing changes. And importantly, bees have displayed strikingly different splicing patterns depending on whether they actually learn or not during training. The most prominent changes occur in the exon 6 cluster, which compared to *Drosophila* has expanded by additional isoforms but also more frequent use alternative 5′ and 3′ splice sites, which is constituent with the more sophisticated neuroanatomy and behavioral performance of bees compared to *Drosophila*. In contrast, exon 4 and 10 clusters in bees have less isoforms. Lower diversity in these two clusters has been suggested to be associated with Dscam’s role as a pathogen receptor in the immune system as flies are extensively exposed to pathogens due to their life in a decomposing environment (Graveley et al., 2004).

*Dscam* diversity has been shown to change in mosquitos towards isoforms that recognize pathogens with higher affinity (Dong et al., 2006). Our finding that *Dscam* splicing changes upon learning further supports that the *Dscam* splicing pattern is not fixed and can change on demand. In fact, levels of Dscam have been found critical to neuronal function in *Drosophila* affecting nerve growth, synaptic targeting and neuronal physiology (Cvetkovska et al., 2013, Lowe et al., 2018, Hernández et al., 2023). Moreover, *Dscam* has also been identified as a target of Fragile X messenger ribonucleoprotein 1 (Fmr1) that together with mRNA modification pathways impacts on local translation important for neuronal functions such as learning and memory (Cvetkovska et al., 2013, Haussmann, 2022, Anreiter et al., 2023).

Consistent with subtle changes in Dscam expression being effective in altering nerve growth, synaptic targeting and neuronal physiology, we find changes in alternative splicing during memory consolidation. In particular, skipping of the entire variable cluster occurs for exon 4 and 10 shortly after training resulting in removal of part of the homophilic interaction domain concomitantly reducing cell adhesion (Wojtowicz et al., 2004).

Dynamic regulation of Dscam expression in the nervous system is somewhat unexpected based on Dscam’s role in mediating axonal bifurcation for exit from axonal tracts or spreading in dendritic fields based on Dscam’s homophilic repulsive properties. Potentially, Dscam’s repulsive homophillic properties could only apply during development of the nervous system and convert to attraction to expand connectivity in mushroom bodies to consolidate memories. In fact, attractive properties of Dscam have been found with lower Dscam concentration (Wojtowicz et al., 2004). Moreover, Dscam has been shown roles in directed wiring of PNS neurons to innervate the ventral nerve cord (Chen et al., 2006). Likewise, whether homophilic binding of Dscam turns into repulsive or attractive cues is also dependent on interacting partners such as Netrins, Robo and Slit (Alavi et al., 2016, Andrews et al., 2008, Liu et al., 2009).

How *Dscam* mutually exclusive alternative splicing is regulated is still poorly understood. Initially, it has been proposed that base-pairing of a constant region at the beginning or end of the cluster with sequences in front of variable exons would direct inclusion of variable exons, but compelling sequence conservation is only present in the exon 6 cluster (Haussmann et al., 2019, Graveley, 2005, Ustaoglu et al., 2019). Our finding that inclusion of alternative exons from the variable clusters changes upon memory consolidation further suggests that there is a mechanism to alter inclusion of specific variables possibly to change neuronal connections for enhancing memory. Given an increasingly aging human population, deciphering the molecular mechanism underlying memory enhancement is of great interest in light of cognitive disfunctions associated with Alzheimer’s, Parkinson and other age-related diseases (Stern and Alberini, 2013, Noyes et al., 2021).

## Materials and Methods

### Honey bees, treatment and behavioral assays

For initial molecular analysis of honey bee Dscam, bees (*Apis mellifera*) were collected from local bee hives in the UK (worker bees were used unless otherwise specified). For developmental gene expression analysis and behavioral experiments, bees were taken from the experimental apiary on the university campus in Toulouse (France), on the morning of each experiment. For behavioral experiments, workers were harnessed in metal tubes following cold-anesthesia leaving access to the head, fed with 5 µl of sucrose solution (50% weight/weight in water) and until needed kept in the dark at room temperature. They were fed in the same way on every morning and evening during the time of each experiment. Learning and memory experiments were done as described (Ustaoglu et al., 2021).

### RNAi, Western analysis recombinant protein expression antibody stainings and imaging

For RNAi knockdown in bees, a *Dscam* DNA template of 820 bp was amplified spanning a cDNA of exons 11-14 with primers AM Dscam T7 RNAi 11F and AM Dscam T7 RNAi 14R containing a T7 promoter on each side and cloned into into pB SK+ tango with Xho and EcoRV according to standard procedures (Soller and White, 2005). A 700 bp fragment for *GFP* was amplified as described (Ustaoglu et al., 2021). Double stranded RNA was generated by *in vitro* transcription with T7 polymerase with the MegaScript kit (Ambion) for 3 h according to the manufacturer’s instructions. After digestion of the template with TurboDNAse (Ambion), dsRNA was phenol/chloroform extracted, ethanol precipitated and taken up in RNAse free water at a concentration of 5 µg/µl. The dsRNA (250 nl) was then injected into the brain through the median ocellus with a Nanoject II microinjector (Drummond).

RNAi efficiency testing for Dscam was done from dissected central brains by Western blotting according to standard protocols as described (Soller and White, 2005) using a polyclonal rat anti-Dscam antibody (1:1000, 358 against a constant part from amino acid 1650 to 2016) (Watson et al., 2005) and infrared dye coupled secondary antibodies (IRDye800CW, LI-COR) were used and detected with an Odyssey infrared imaging system (LI-COR). Tubulin was detected with a mouse anti-alpha tubulin antibody (1:10,000, clone DM1A, SIGMA). Quantification of Western blots was done with Quantity ONE 4.6.8 (BioRad) according to the manufacturer’s instructions.

### RT-PCR and analysis of alternative splicing

The sequence of oligonucleotides used in this study are listed in Supplementary Table 1. RNA extraction from eggs or dissected central brains and RT-PCR was done as described (Koushika et al., 1999). For RT, AM GSP exon 13 RT1 was used at concentration of 10 nM spiked into oligo dT (10 µM) (Haussmann et al., 2011).

For high resolution analysis of *Dscam* alternative splicing primers AM Dscam 3F1MH and AM Dscam 5R1 were used to amplify exon 4 variables, and AM Dscam 9F2 and AM Dscam 11R2 for exon 10 variables from cDNA by PCR with Taq according to the manufacturer’s instructions for 40 cycles as described (Soller and White, 2005). One of the primers was labeled using gamma^32^P-ATP (NEN). PCR products for exon 4 variables were digested with a combination of restriction enzymes Sau3AI, HaeIII, and MboI, and PCR products for exon 10 variables with a combination of restriction enzymes RsaI, PvuII and BsaJI. Then, PCR products were separated on 5 % sequencing type denaturing polyacrylamide gels. Polyacrylamide gels were dried, exposed to phosphoimager screens (BioRad) and quantified with QuantityOne (BioRad).

### Amplicon sequencing and sequence analysis

For amplicon sequencing, primers in constant exons flanking the variable parts were used for amplification from cDNA generated from dissected central brains as described above. For amplification of variable exon 6 from three groups of three individual bees AM Dscam 5F1A, B and C were used for the different groups and AM Dscam 7R1A, B and C were used for the three individual bees. Amplicons were then quantified on ethidium bromide stained agarose gels against a DNA marker (100 bp ladder, NEB), pooled and then Illumina sequenced (paired end, read-length 126) by GATC (Eurofins, Germany). Sequences were demultiplexed according to barcodes for exon 6 variables or by sequence cluster identity for exon 4 and 10 variables and barcodes removed using an in-house script. The demultiplexed data was mapped to the Bee reference genome (version Amel_HAv3.1) using STAR (STAR version 272B, parameters: --alignIntronMin 20 --alignIntronMax 200000 --alignMatesGapMax 200000 --twopassMode Basic). Junction read coverage matching Dscam (AY686596) (Graveley et al., 2004) have been used to compute inclusion levels of splicing events, following which junction reads were manually checked.

### Statistics and reproducibility

Multiple planned pairwise comparisons of expression levels were done by ANOVA followed by Fisher’s protected least significance difference post-hoc test using Prism. To compare proportions of conditioned responses between groups, a repeated-measure analysis of variance (ANOVA) was run for the acquisition data (one factor, *treatment*, with *trial* as the repeated measure) (Mota and Giurfa, 2010). Comparisons of responses at each memory test were run using McNemar and Fischer’s exact tests, for intra- and inter-group comparisons respectively.

## Acknowledgments

We thank the Winterbourne garden (Birmingham), Jane and Lucie Hotier (Toulouse) for providing bees, Dietmar Schmucker for a rabbit anti-Dscam antibodies, Anna Lassota for help with stainings and imaging, Reinhard Stöger for discussions and comments on the manuscript. For this work we acknowledge funding from the Sukran Sinan Fund, the Genetics Society, the Biochemical Society, MRC DTP and BBSRC. JMD acknowledges funding from the CNRS and Université Paul Sabatier.

## Authors’ contribution

PU, IUH, TD and MS performed molecular biology experiments, PU, AR and JMD performed behavioral experiments, and PU and TD performed antibody stainings, Western blots and imaging. JMD designed behavioral experiments. DWJM and RA analysed sequences. MS conceived the project and wrote the original draft of the manuscript. JMD, IUH and all other authors reviewed and edited. MS and JMD supervised and acquired funding.

## Ethics statement

Ethical review and approval was not required for this study because this study was conducted with an invertebrate model – honey bee (*Apis mellifera*). Experiments with invertebrates are not regulated by law.

## Data availability

All data are available in the main text or the supplementary material (Supplementary Data 1). Reagents are available upon reasonable request from the corresponding author. Amplicon-Seq data have been deposited at GEO (GSE244051, reviewer token: anwxcquopdgxpav).

---

**Suppl. Fig. 1.**
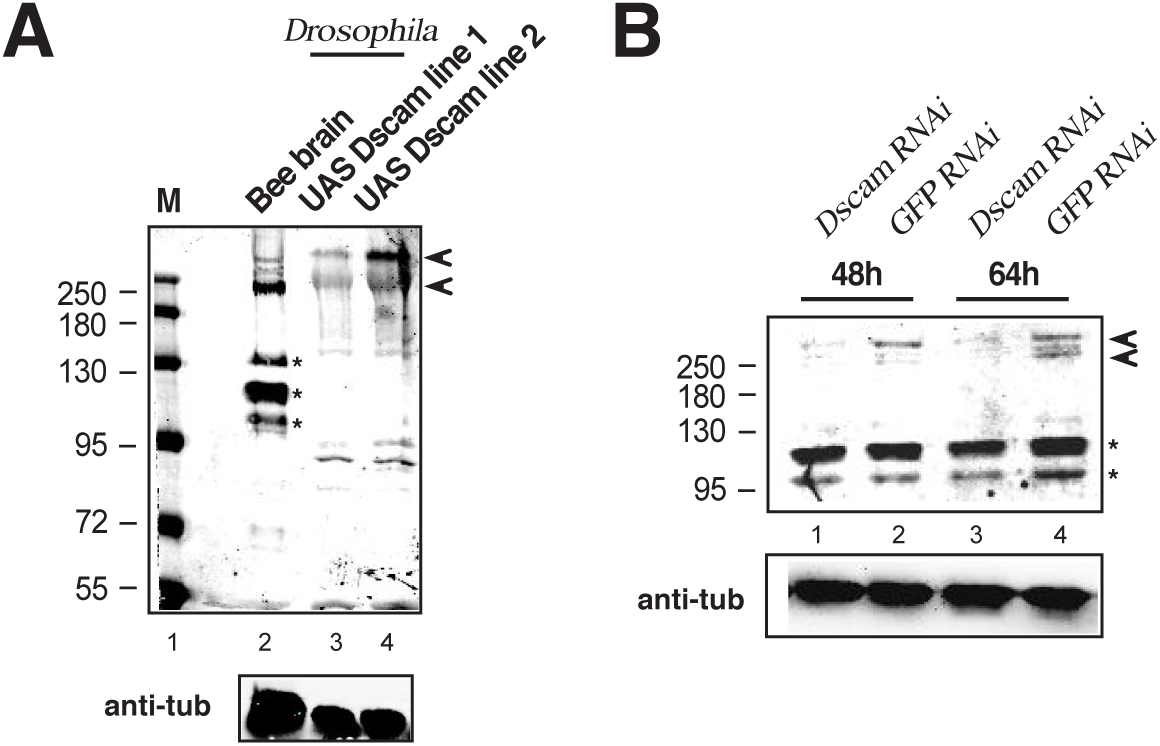
*Drosophila* anti-Dscam recognizes honey bee Dscam. **A** A rabbit polyclonal anti-serum raised against *Drosophila* Dscam recognizes honey bee Dscam on Western blots at expected sizes of 222 and 270 kDa (arrowheads). Extracts of bee brains (lane 2) or head and thorax from *Drosophila* expressing *UAS Dscam* with *daughterlessGAL4* (lanes 3 and 4) were separated on a 8% SDS-gel. Note that the anti-serum recognizes unspecific proteins of 100, 110 and 130 kDa in bees (asterisks). Loading is shown by probing the same blot with anti-tubukin antibodies (bottom). M: Molecular weight marker. **B** Western blot of bee brains 48 and 64 h after injection of *Dscam* or *GFP* dsRNA for RNAi knock-down. Note that Dscam levels are reduced after RNAi by injection of Dscam dsRNA (lanes 1 and 3), but not the unspecific proteins of 100, 110 and 130 kDa (asterisks). Loading is shown by probing the same blot with anti-tubulin antibodies (bottom). Molecular weight markers are indicated on the left.

**Suppl. Fig. 2.**
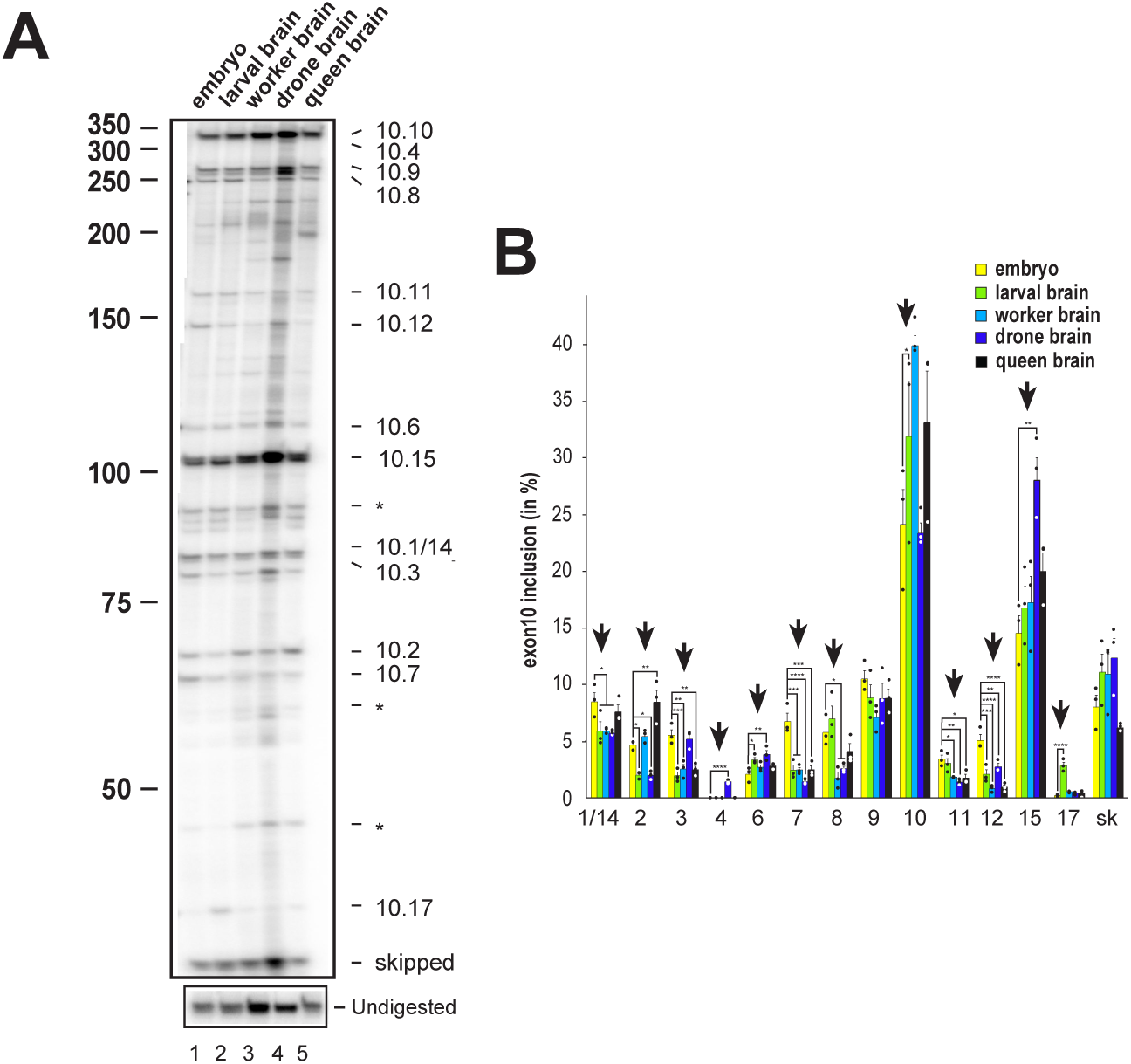
*Apis mellifera Dsacm* exon 10 alternative splicing during bee development and between casts. **A** Denaturing polyacrylamide gels showing the splicing pattern of *Dscam* exon 10 isoform variables on top by digestion of a ^32^P labeled RT-PCR product with a combination of *Rsa*I, *Pvu*II and *BsaJ*I restriction enzymes in embryos (line1), larval brains (line 2), worker brains (line 3), drone brains (line 4) and queen brains (line 5) and undigested control at the bottom. sk: skipping of all exon 10 variables. Molecular weight markers are indicated on the left. **B** Quantification of inclusion levels of individual exons are shown as means with standard error from three experiments for embryos, larval brains, worker brains, drone brains and queen brains (*: p<0.05; **: p<0.01; ***: p<0.001; ****: p<0.0001).

